# Live imaging YAP signaling in mouse embryo development

**DOI:** 10.1101/2021.08.11.456026

**Authors:** Bin Gu, Brian Bradshaw, Min Zhu, Yu Sun, Sevan Hopyan, Janet Rossant

## Abstract

YAP protein is a critical regulator of mammalian embryonic development. By generating a near-infrared fusion YAP reporter mouse line, we have achieved high-resolution live imaging of YAP localization during mouse embryonic development. We have validated the reporter by demonstrating its predicted responses to blocking LATs kinase activity or blocking cell polarity. By time lapse imaging preimplantation embryos, we revealed a mitotic reset behavior of YAP nuclear localization. We also demonstrated deep tissue live imaging in post-implantation embryos and revealed an intriguing nuclear YAP pattern in migrating cells. The YAP fusion reporter mice and imaging methods will open new opportunities for understanding dynamic YAP signaling *in vivo* in many different situations.

## Background

*Yes associated protein 1 (Yap1)*-commonly referred to as YAP, is a signaling protein that serves as a hub of biochemical and mechanical sensing^1–3^. YAP plays crucial regulatory roles in enormously diverse processes in development and disease, from the very beginning of life, such as mammalian preimplantation development^4^, to the very late stage of disease, such as cancer metastasis^5^. Real-time tracking of the dynamic YAP signaling status *in vivo* would offer crucial insights into the fundamental regulatory mechanisms of mammalian development and disease.

YAP is a transcriptional co-activator for the TEA domain (TEAD) family transcription factors^6^. In response to numerous signals, YAP shifts its sub-cellular distribution to the nucleus where it associates with TEAD proteins to activate gene expression. Thus, monitoring the nuclear/cytoplasmic localization of YAP is the most broadly applied method to evaluate YAP activity^6,7^. Currently, this is primarily achieved by immunostaining of fixed embryos or tissue sections, precluding the acquisition of dynamic information. Although live imaging has been achieved in Drosophila and human cell lines and revealed intriguing YAP behaviours^8,9^, no appropriate tools exist to date for live imaging endogenous YAP protein in mammals *in vivo*. Here we report a YAP fusion reporter mouse line that allows in vivo live imaging of YAP behaviour.

## Methods

### Designs of guide RNAs and knock-in repair donors

Guide RNAs target spanning the Stop TAG codon: TCACGTGGTTATAGAGCTGC**AGG** were selected using the CRISPOR algorism (http://crispor.tefor.net) based on specificity scores^10^. The chemically modified single guide RNAs (sgRNAs) with the sequence of UUGCGCGGGCUCCAUGGCUG were synthesized by Synthego Inc. The repair donor for YAP-emiRFP670 reporter were designed as illustrated in Extended Figure 1A. The emiRFP670 and the linker coding sequences in proper orientation were flanked by long homology arms (813bp 5’ arm and 717bp 3’arm) on each side and replacing the stop codon of the *Yap1* gene. The Donor DNA sequence was synthesized by Epoch Life Science (http://www.epochlifescience.com) and cloned into a PBSK plasmid backbone.

### Generating KI reporter mouse lines by 2C-HR-CRISPR

The KI reporter mouse line was generated following our published protocol using 2C-HR-CRISPR on the CD1 background^11,12^. Briefly, Cas9 monomeric streptavidin (mSA) mRNAs were produced by in vitro transcription using the mMessage mMachine SP6 Kit (Thermo Scientific). PCR templates were generated by PCR reaction with biotinylated primers using high fidelity ClonAmp HiFi PCR mix (Takara Inc.). A mixture of Cas9-mSA mRNA (75ng/μl), sgRNA (50ng/μl), and biotinylated repair donor (20ng/μl) was microinjected into the cytoplasm of 2-cell stage mouse embryos and transferred the same day to pseudo-pregnant females. Founder pups were obtained from the pregnancies. We established founder mice with the correct insertion at high efficiency (2/2 live born pups). A founder was outcrossed five generations to wildtype CD1 mice to breed out any potential off-target mutations introduced by CRISPR-Cas9 and then bred to homozygosity at N6 generation. The mouse line was then maintained by homozygous breeding. In the early generations of homozygous breeding, a cataract phenotype was observed in some mice. This phenotype was then removed by selectively breeding YAP-emiRFP670 homozygous reporter mice without such a phenotype. The homozygous mice are otherwise healthy and fertile without apparent phenotype.

For generating YAP-emiRFP670/Cdx2-eGFP embryos, the YAP-emiRFP670 mouse line was bred to the Cdx2-eGFP mouse line to generate double homozygous mice^13^. Embryos were then collected from the double homozygous mouse line for imaging.

All animal work was carried out following the Canadian Council on Animal Care Guidelines for Use of Animals in Research and Laboratory Animal Care under protocols approved by the Centre for Phenogenomics Animal Care Committee (20-0026H).

### Genotyping and genetic quality controls

Founder mice were genotyped by PCR amplification with primers spanning homology arms using the following primers: 5’ arm gtF: GTTCTAAGGTAGACACTGTGTGCTTCAGTT and 5’arm gtR: TCATGTTCGCAGGTCAAGAGGTCA; 3’arm gtF: CTGGTTGTCTGTCACCATTATCTGC and 3’arm gtR: AACACCTGCAATTGCTCCAACC. Founder mice were out-crossed to CD1 mice to generate N1 mice. The N1 mice were genotyped by PCR. Additionally, genomic regions spanning the targeting cassette and 3’ and 5’ homology arms were Sanger-sequenced to validate correct targeting and insertion copy number was evaluated by droplet digital PCR (performed by the Centre for Applied Genomics at the Research Institute of The Hospital for Sick Children, Toronto). Heterozygous N1 mice have only one insertion copy, demonstrating single copy insertion.

### Embryo isolation, culture, and treatments

Preimplantation embryos were isolated from superovulated females that were mated with males, both homozygous for the C-YAP reporter. Embryos were isolated from oviducts or uterus at appropriate stages for each experiment – E0.5 for zygote, E1.5 for 2-cell embryos, E2.5 for 8-cell embryos and E3.5 for blastocysts. Embryos were flushed with M2 medium. They were then cultured in small drops of KSOM-AA medium under mineral oil at 37°C, with 6% CO2 for specified times. For the Rock inhibitor treatment experiment, 8-cell stage embryos were cultured in KSOM-AA medium supplemented with 20μM Y-27632.

### Time Lapse Live imaging preimplantation embryos

For imaging the YAP dynamics during the formation of 16-cell embryos, embryos were flushed from the oviduct at 4-cell or 8-cell stage and cultured in a 3ul drop of KSOM-AA medium under mineral oil on a MatTek glass bottom dish. Live imaging was performed at 20mins/frame for 24-36 hours. For the ROCKi inhibitor treatment, embryos were flushed from the oviduct at 8-cell stage and cultured in a 3ul drop of KSOM-AA medium with 20μM Y-27632 under mineral oil on a MatTek glass-bottom dish. Live imaging was performed every 90 mins for 24 hours.

### dnLATS2 mRNA injection

Homozygous YAP-emiRFP670 embryos were collected at 2-cell stage. mRNAs were microinjected into one of the two blastomeres as previously described^7,14^. For control experiments, embryos were injected with H2B-RFP mRNAs at 300ng/μl. Treated embryos were injected with dnLats2 mRNA at 1000ng/μl plus H2B-RFP mRNA at 300ng/μl. The embryos were then cultured to the early blastocyst stage for immunofluorescent analysis.

### Whole-mount immunofluorescent staining of embryos

Whole mount immunofluorescent staining of embryos was performed as previously described^15^. Briefly, embryos were fixed in 4% paraformaldehyde at room temperature for 15 minutes, washed once in PBS containing 0.1% Triton X-100 (PBS-T), permeabilized for 15 minutes in PBS 0.5% Triton X-100 and then blocked in PBS-T with 2% BSA (Sigma) and 5% normal donkey serum (Jackson ImmunoResearch Laboratories) at room temperature for 2 hours, or overnight at 4°C. Primary and secondary antibodies were diluted in blocking solution, staining was performed at room temperature for ~2 hours or overnight at 4°C. Washes after primary and secondary antibodies were done three times in PBS-T. Nuclear staining was performed using Hoechst 33258 (Thermo scientific) at a concentration of 10μg/ml for 20 minutes at room temperature. Embryos were mounted in PBS in wells made with Secure Seal spacers (Molecular ProbesTM, Thermo Fisher Scientific) and placed between two cover glasses for imaging. Primary antibodies: Goat anti-tdTomato (Biorbyt orb182397 Lot AR2150) at 1:200 dilution, Rabbit anti YAP (Cell Signalling Technology (D8H1X) XP Ref 11/2018 Lot4) at 1:100 dilution and Mouse anti Cdx2(Biogenex MU392A-UC Lot MU392A0714) at 1:100 dilution. Secondary antibodies all from Themo Scientific and at 1:400 dilution: Donkey-anti-goat IgG AF 546 (A11056 Lot 1714714), Donkey anti Rabbit IgG AF647(A31573 Lot 1693297) and Donkey anti mouse IgG AF488 (A21202 Lot 1741782)

### Confocal imaging of preimplantation embryos

Both live and immunostained images of preimplantation embryos were acquired using a Zeiss Axiovert 200 inverted microscope equipped with a Hamamatsu C9100-13 EM-CCD camera, a Quorum spinning disk confocal scan head, and Volocity acquisition software version 6.3.1. Single plane images or Z-stacks (at 1μm intervals) were acquired with a 40x air (NA=0.6) or a 20x air (NA=0.7) objective. Images were analyzed using Volocity software. Live imaging was performed in an environment controller (Chamlide, Live Cell Instrument, Seoul, South Korea) on the same microscope.

Time-lapse imaging was performed on the same microscope equipped with an environment controller (Chamlide, Live Cell Instrument, Seoul, South Korea). Embryos were placed in a glass-bottom dish (MatTek) in KSOM-AA covered with mineral oil. A 20x air (NA=0.7) objective lens was used. Images were acquired at 1-3μm Z intervals with time-lapse settings as indicated in figure legends.

### Image quantification analysis for preimplantation embryos

Preimplantation images were visualized using the Volocity 6.3 software. The live images of preimplantation embryos were traced manually by carefully inspecting the movie and tracking the cell nucleus marked by the live DNA dye and recording the presence or absence of the C-YAP reporter. For quantifying fluorescent intensity, images were exported as TIFF files and measured using the Region of interest (ROI) function in ImageJ 1.53a software. The average fluorescent intensity of each nuclear ROI was measured and subtracted against a general background fluorescent intensity in the corresponding image.

### Lightsheet imaging of post-implantation embryos

Three-dimensional static live imaging was performed on a Zeiss Lightsheet Z.1 lightsheet microscope. Embryos were suspended in a solution of DMEM without phenol red containing 12.5% filtered rat serum and 1% low-melt agarose (Invitrogen) in a glass capillary tube. Once the agarose solidified, the capillary was submerged into an imaging chamber containing DMEM without phenol red, and the agarose plug was partially extruded from the glass capillary tube until the portion containing the embryo was completely outside of the capillary. The temperature of the imaging chamber was maintained at 37° C with 5% CO2. Images were acquired using a 20× objective with dual-side illumination in a multi-view mode (4 evenly spaced views spanning 360° for E8.0 embryos imaging) or tile scanning mode (for E8.5 and E9.5 embryos imaging). The light sheet images were processed using Zen (Zeiss), Arivis Vision4D (Arivis), Imaris (Bitplane) and ImageJ.

Time lapse imaging was performed similarly to the static live imaging with the following modifications: 1) 2% fluorescent beads (1:1,000, diameter: 2 μm; Sigma-Aldrich) were added to the low melting point agarose for drift-compensation. 2) Images were acquired for 3 hours with 5 min. intervals.

### In vivo drift-compensated cell tracking and mean squared displacement calculation

The light sheet time-lapse image was first rendered in Imaris (Bitplane). Nuclear YAP positive cells were determined by mean thresholding of fluorescence intensity. For each embryo, the cut-off intensity was set to be fifty percent of its maximum intensity. Small bright spots of YAP (diameter < 7 μm) due to local chromatin condensation were excluded from nuclear YAP positive cell identification. The positions of nuclear YAP positive cells were tracked over time using an autoregressive motion algorithm. The tracking data were then imported into MATLAB (MathWorks) for drift compensation using a program reported before^16^.

Mean squared displacement (MSD) is an unbiased metric to evaluate cell migration^17^. For an arbitrary trajectory, the MSD and time delay follows a power-law relation, with power of 0 representing the random walk motion, and power of 2 representing the straight-line motion. On a log–log plot, these two cases translate into a line with slope 0 (i.e., a horizontal line) for the random walk motion, and a line of slope 2 for the straight-line motion. The mean MSD of nuclear YAP positive cells’ migration trajectories are characterized by a line of slope 1.433 as shown in Fig. 4C which deviates significantly from the random walk motion suggesting a persistent cell migration^18^.

### Statistical analysis

Statistics on numerical data were performed using the Prism 9 software(GraphPad Software, LLC.). For intergroup comparison, the data were first subjected to the D’Agostino & Pearson normality test. Datasets that conform to a normal distribution were then subjected to the unpaired Student’s-t test, and the ones that did not conform to a normal distribution were then subjected to the Mann Whitney test. 2-tailed analysis were used in all cases. Exact p-values were presented in the figures. For the one group t-test for the proportion of relative position of cell division pairs at 16-cell stage, a null hypothesis of the proportion= 50% was tested against and resulted in a p-value (p=0.0381).

### Results and discussion

We set out to engineer a knock-in (KI) fusion reporter of *Yap* in mice (Extended Figure 1). There have been several previous attempts to generate endogenous *Yap* KI reporters, including by ourselves, without success. The function of mammalian YAP seems to be easily disrupted by fusion tags. We tested different combinations of tagging position and linker sequences and finally successfully generated a healthy reporter mouse line by C-terminal tagging with a long (30 amino acid) flexible linker (Extended Figure 1A). Other designs, such as N-terminal tagging with the same linker led to embryo death at embryonic days 8.5 (E8.5), similar to Yap knockout embryos, suggesting functional interference^19^. To allow good light penetration for imaging deep tissue layers in postimplantation mammalian embryos and other tissues, we chose to use a bright near-infrared (NIR) fluorescent protein (FP), enhanced miRFP670(emiRFP670), as the fluorescent indicator^20^. We performed extensive quality control to confirm single-copy insertion of the fusion reporter with the correct sequence, as detailed in the methods section and extended Figure 1B and our published protocol^11,21^. The homozygous mice are healthy and fertile. The mouse line was maintained in the homozygous state after outcrossing for 5 generations to ensure no carry-over of any possible off-target alterations. We validated the reporter in mouse preimplantation embryos where the YAP distribution and its responses to various interventions are well characterized. The nuclear/cytoplasmic distribution of YAP from 8-cell stage onward is well established from previous research: At 8-cell stage, all blastomeres have nuclear localized YAP. From 16-cell stage to early-mid blastocyst stage (E3.5), nuclear YAP is restricted to the outer cells that will give rise to the trophectoderm^7^. By the expanded blastocyst stage (E4.5), some epiblast cells will start to present nuclear YAP status^22^. Our live imaging reproduced this pattern (Figure 1A). Before the 8-cell stage, the localization of YAP distribution is more debatable. Our live imaging showed that YAP was evenly distributed in the cytoplasm and nucleus of blastomeres until the late 4-cell stage, at which point YAP begins to be more restricted to the nucleus (Figure 1A). We further validated the colocalization of the C-YAP reporter with the endogenous YAP protein and with the expression of CDX2 protein in the TE of blastocysts by immunofluorescence (Figure 1B).

**Figure 1:**
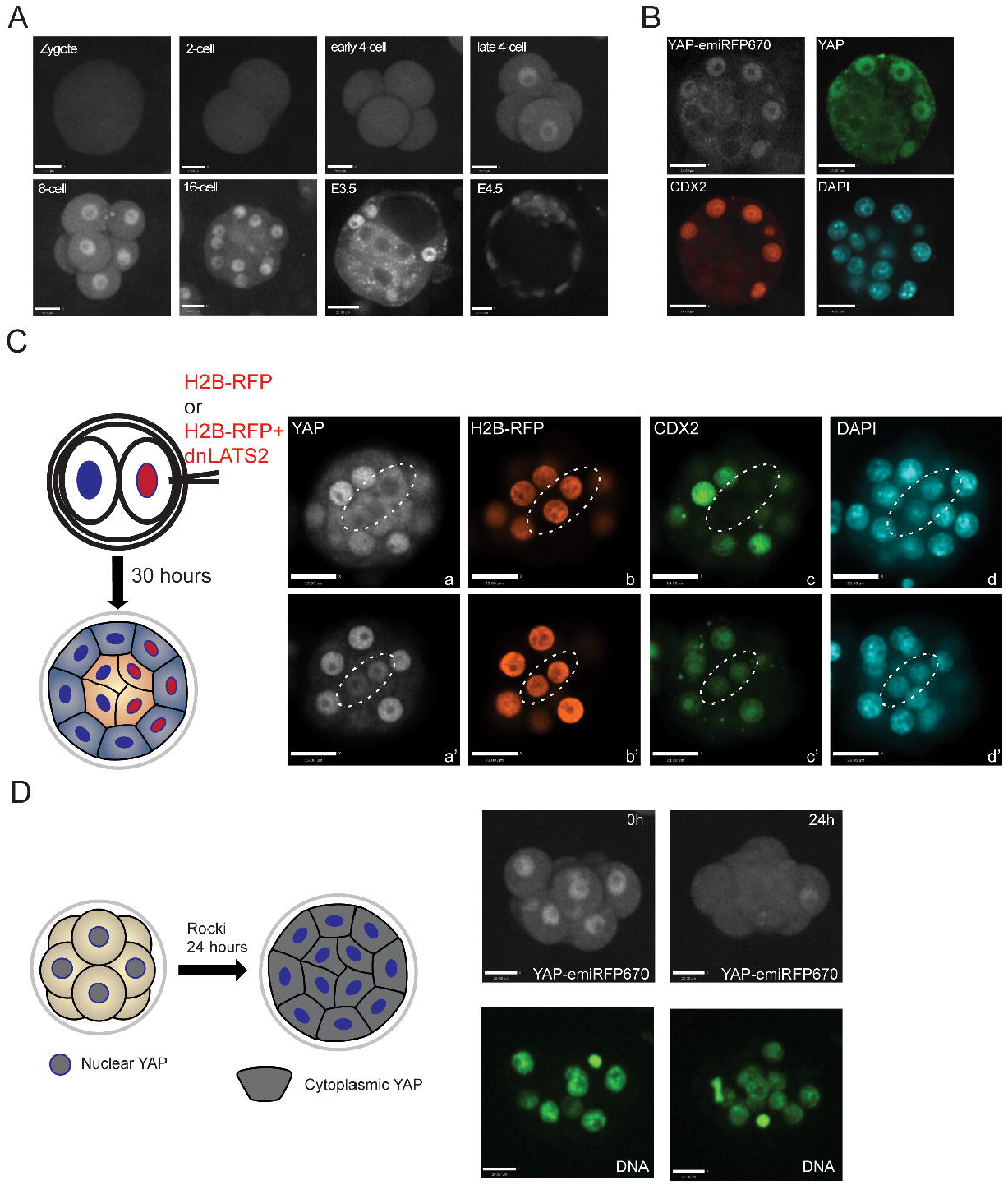
Characterization and validation of C-YAP reporter in preimplantation embryos: A. Live imaging of C-YAP localization in preimplantation embryos. B. Immunofluorescence images of the C-YAP reporter (emiRFP670 fluorescence), endogenous YAP protein (Immunofluorescence) and CDX2 protein (Immunofluorescence) in a mouse blastocyst, showing perfect colocalization of YAP and CDX2 in the embryo. C. Manipulation of YAP-emiRFP670 localization by expressing a dominant-negative LATS2. Left: A schematic for mRNA injection experiment. mRNAs of H2B-RFP (control group) or H2B-RFP+dnLATS2(experimental group) were injected into one of the two cells of a 2-cell stage mouse embryo and then cultured to early blastocyst stage for analysis. Right upper panel, a control embryo shows that the expression of H2B-RFP did not cause nuclear YAP localization and CDX2 expression. Right bottom panel, a dnLATS2 injected embryo showed nuclear YAP and CDX2 expression in inside cells. Cells of interest were circled by dotted lines. D. Manipulation of YAP-emiRFP670 localization by treating with the ROCK inhibitor Y27632. Left: A schematic for the treatment experiment. Right, Snapshots from live imaging series Movie 1 showing ROCK inhibitor treatment leads to cytoplasmic Yap localization in embryos.

We then investigated whether the C-YAP reporter protein can respond appropriately to exogenous signals. In mouse early embryos, YAP localization is controlled by HIPPO and polarity signaling pathways. From the HIPPO signaling pathway, LATS1/2 kinase is responsible for the phosphorylation of YAP, which leads to its sequestration in the cytoplasm^7^. Overexpressing a kinase-dead LAT kinase (dnLATS2) can promote the nuclear localization of YAP and the expression of CDX2 even in inside cells^7^. This result was replicated when we injected a dnLATS2 mRNA into homozygous C-YAP embryos (Figure 1C). As for polarity, it has been shown that the inhibition of the Rho-associated protein kinase (ROCK) kinase using a small molecular inhibitor Y27632 resulted in universal cytoplasmic YAP localization, and as a result, failure to establish the TE^23^. We cultured C-YAP reporter embryos from 8-16 cell stages in Y27632 and live imaged them. As shown in Figure 1D and movie 1, ROCK inhibition indeed resulted in a cytoplasmic YAP localization in all cells (Figure 1D).

We then used the validated reporter mouse line to analyze the behaviour of YAP during embryonic development in real-time. A key event that YAP regulates during preimplantation development is the initiation of inner cell mass (ICM)-TE segregation at the 16-cell stage. When 8-cell embryos transition to 16-cell embryos through cell division, YAP becomes localized in the nucleus of blastomeres located on the surface of the embryos (outside cells) and excluded from the nucleus in inside cells^7^. The presence of nuclear YAP drives the expression of the TE-specific transcription factor CDX2 in outside cells, initiating TE lineage differentiation. In contrast, the enclosed blastomeres with cytoplasmic localized YAP initiate *Sox2* expression and ICM differentiation^24^. However, the exact process that leads up to this asymmetric YAP signaling status is still debatable. Some models suggest a fixed determination of YAP localization at the time of generation of inside and outside 16-cell blastomeres, while other recent studies suggesting a more dynamic process^25,26^. Direct observation of the YAP dynamics through the 8-16 cell stage is the best way to resolve these models.

We time-lapse imaged homozygous C-YAP reporter embryos every 20 mins at the transition between the 8-cell to 16-cell stage for a period of 20 hours. Cell nuclei were marked by a live DNA dye (Movies 2-3, with additional movie from 4-cell stage-Movie 4). To rule out the possibility of the imaging process affecting normal development, we further cultured the embryos to E4.5 and validated a high blastocyst formation rate (8/9) (Extended Figure 2A). The movies revealed profound dynamic movements of blastomeres in 8-16 cell stage embryos. Many cells changed their relative outside-inside position from where they were localized right after the cell division that generates them (Movies 2 and 3). We discovered an intriguing mitotic reset behavior pattern of YAP during this transition through closer inspection and tracking cells from the movies. When an 8-cell blastomere divides to form two daughter cells - 16-cell blastomeres- both always show nuclear localization of YAP, regardless of the direction of the cell division axis (Figure 2A). It took roughly 100-300 mins for the two daughter cells to adopt their final position in the embryo (Figure 2A). The final position that a cell adopted determined its YAP distribution- when a cell adopted an outside position, it presented nuclear YAP, whereas when a cell adopted an inside position, it presented cytoplasmic YAP (Examples in Figure 2A and quantifications in Extended Figure 2B and C). There was an almost equal chance for two 16-cell blastomeres from a single 8-cell blastomere to adopt one of the two relative positions – outside-outside or outside-inside (Extended Figure 2D). A previous report has suggested that a significant proportion of inside cells at the 16-cell stage are the result of cell movement rather than oriented cell division, consistent with our observations^26^. The YAP-emiRFP670 reporter allowed us to relate this dynamic cell movement to a dynamic regulation of YAP signaling. Interestingly, the movie of embryos treated with ROCK inhibitor (Movie 1 and 5 and Figure 2B) showed that, although ROCKi eventually inhibited YAP nuclear localization in all 16-cell blastomeres, the initial YAP nuclear localization right after cell division was not affected, suggesting a differential involvement of ROCK-related processes, such as polarity and mechanical tension, in regulating these two distinct YAP nuclear localization processes. How this dynamic behavior is controlled remains an open question, which can now be addressed by live imaging the YAP reporter model in combination with additional KI reporters tracking polarity and mechanical tension.

**Figure 2:**
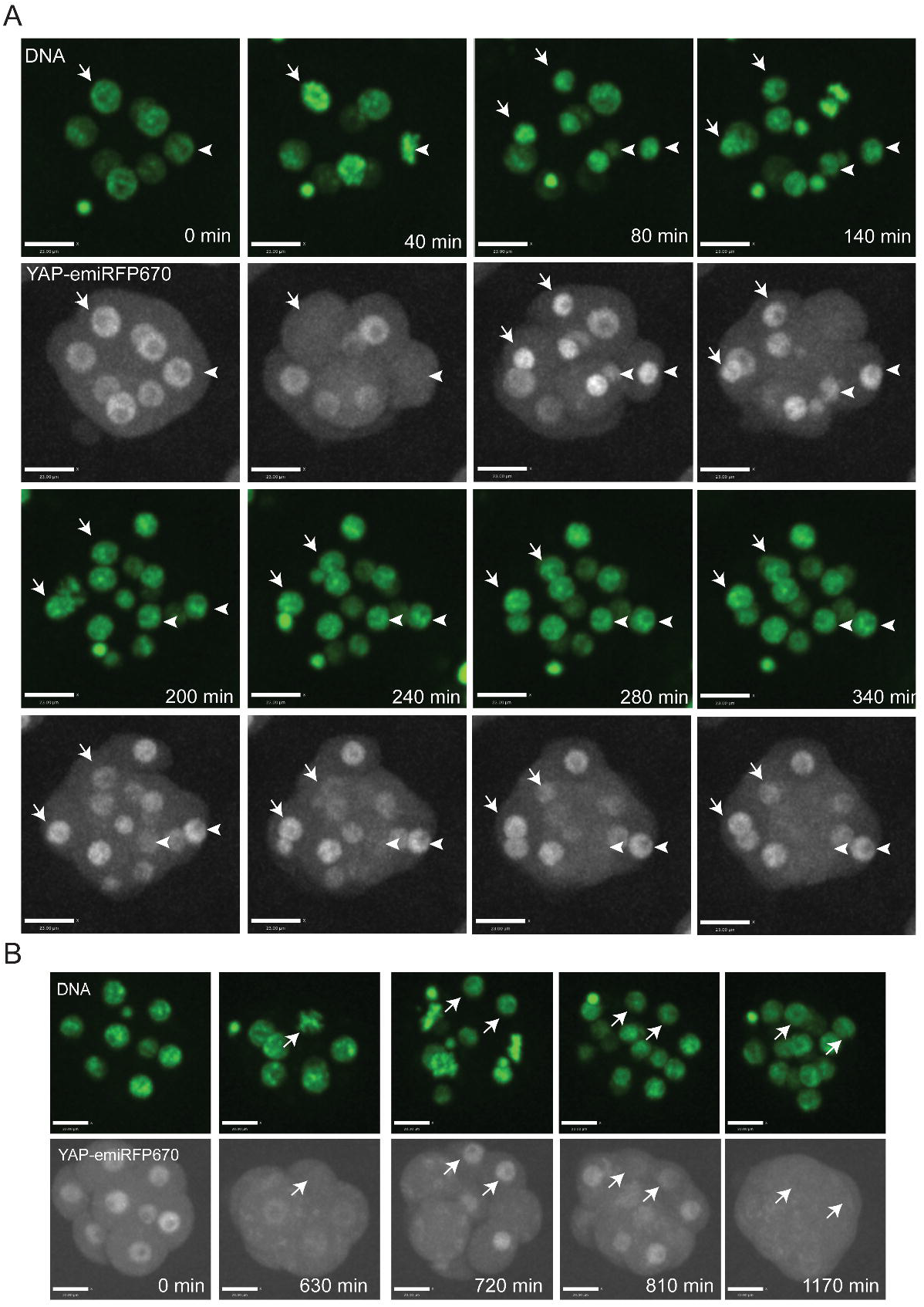
Dynamic YAP localization in 8-16 cell mouse embryos: A. Snapshots from live imaging series (Movie 3) as examples of YAP behavior in mitotic pairs. For the pair annotated by arrows, both cells presented nuclear YAP after the cell division at 80 mins, and both of them located on the outside of the embryo and maintained nuclear YAP at 340 min. For the pair annotated by arrowheads, both cells presented nuclear YAP after the cell division at 80 min. Subsequently one of them was located on the outside of the embryo and maintained nuclear YAP while the other moved to the inside and presented cytoplasmic YAP at 340 min. (Quantitative data in extended figure 3B-D). B. Snapshots from live imaging series (Movie 4) with C-YAP embryos treated with the ROCK inhibitor Y-27632. During the 8-cell stage, 0 min to 630min, ROCK inhibition resulted in primarily cytoplasmic localization of Yap. After cell division, as demonstrated by the cell pair marked by arrows at 720min, all cells showed a transient nuclear localization of Yap. Then all cells gradually reversed to a cytoplasmic YAP localization status over time (810 min and 1170 min).

Preimplantation embryos are small and transparent. To demonstrate the broader application of the YAP reporter in more challenging samples, we live-imaged the YAP-emiRFP670 embryos at E8.0 (before turning, Movie 6), E8.5 (after turning, Movie 7), and E9.5 (Movie 8) using light-sheet imaging technology. We achieved high resolution imaging of multiple tissues including those located in deep tissue layers (up to 200μm), such as the heart tube (Figure 3 and z stack in movie 9). In most regions, tissues consisted primarily of cells with cytoplasmic YAP, with only small populations of cells with strong nuclear signals (Figure 3 B-D, examples marked by arrows). In the heart tube, on the other hand, most cells showed nuclear YAP which may be consistent with the mechanical load these cells are subjected to and suggests a crucial role for YAP in heart development^27,28^. All the raw imaging data of the subcellular distribution of YAP in postimplantation embryos will be deposited on open access databases and will serve as a rich resource for studying YAP signaling in mouse embryos.

**Figure 3:**
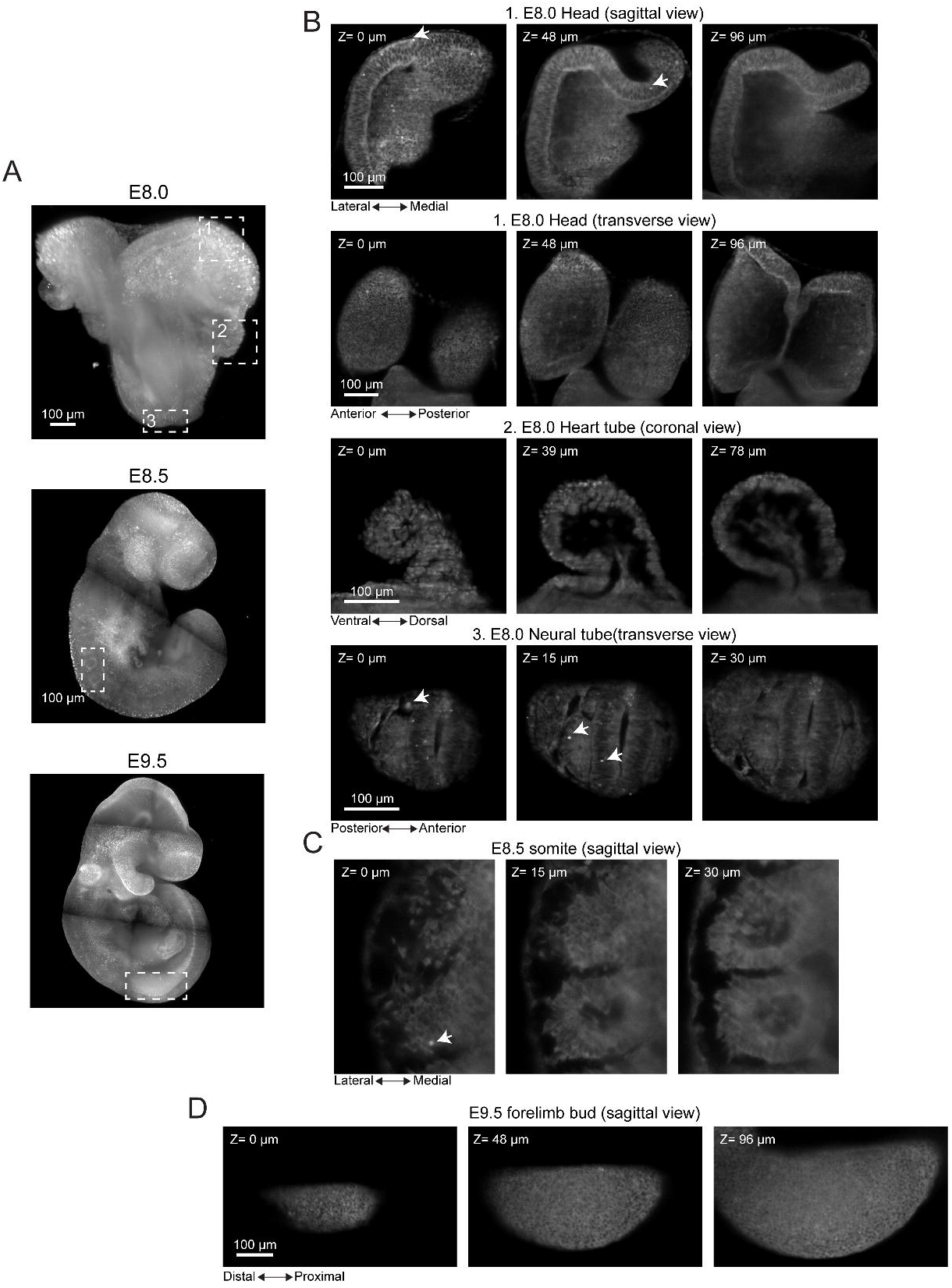
Live imaging of Yap in early organogenesis stage embryos: A. Tile scan images of C-YAP embryos at three developmental stages: E8.0 (before turning), E8.5 (after turning), and E9.5. B. Zoom-in views of square marked regions in the E8.0 embryo: 1. Head fold, 2. Heart tube and 3. Neural tube. C. Zoom-in views of the somite region in the E8.5 embryo (square in image). D. Zoom in views of the limb bud region in the E9.5 embryo (square in image). Arrows in B-D mark examples of cells with strong nuclear Yap.

To reveal the dynamic behavior of cells with active YAP signaling, we time lapse imaged E8.5 embryos every 5 mins for 3 hours (Movies 10). This movie revealed a population of cells with strong nuclear YAP signal migrating within the head region (Figure 4A and movie 10). We conducted tracking of the time dependent motion of these nuclear YAP cells^16^. The persistence of these cellular motions was tested using Mean squared displacement (MSD) from 1962 tracks from three embryos and showed a strong deviation from a random walk model, suggesting persistent cell migration (Figure 4B and C). The identity of these cells and the mechanisms by which YAP regulate directional cell migration in embryos warrants further investigation. These data demonstrate that the C-YAP reporter can have broad imaging applications in challenging samples such as late-stage embryos and tissues.

**Figure 4.**
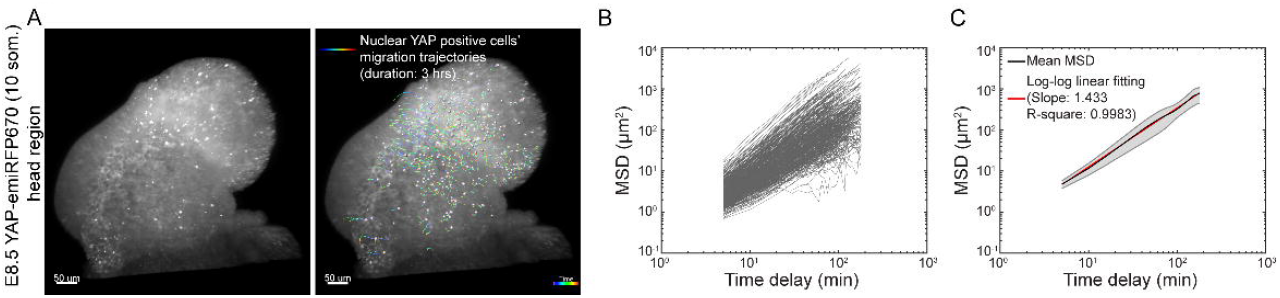
Migrating cells A. Migration trajectories of nuclear YAP positive cells within the head region of a E8.5 YAP-emiRFP670 (10 som.). Left image shows an embryo with bright nuclear YAP cells at the beginning of the time lapse imaging. Right image depicts the migration tracks of these cells over three hours. Colormap represents the time points of individual cell migration trajectories over 3 hours imaging with blue and red being 0 and 3 hours, respectively. B. MSD (mean squared displacement) of nuclear YAP positive cells’ migration trajectories shown in A (520 tracks). C. Mean MSD (black line) of nuclear YAP positive cells’ migration trajectories at E8.5 (10~11 som.) head regions (1962 tracks from three embryos). Log-log linear fitting (red line) yields a slope of 1.433 indicating a persistent cell migration motion (Methods). Gray color marks the standard error of mean.

### Open up

The YAP-emiRFP670 mouse line will open up broad opportunities in biomedical research. For example, existing studies in early embryos suggest that at the very early totipotent stages of development (zygote to pre-compaction 8-cell stage), YAP primarily serves to open up the zygotic genome^29,30^, and then it transforms to a lineage determinant around the 16-cell stage^7^. We conducted a time lapse imaging series with YAP-emiRFP670 and a trophectoderm reporter Cdx2-eGFP (Movie 11-13). The movie revealed very little correlation between the YAP nuclear localization and Cdx2 expression up to the late 16-cell stage. In addition, although ROCK inhibitor treatment caused the cytoplasmic localization of YAP at late 16-cell stage, the CDX2-eGFP persisted (Movie 14). These data suggested a more complex relationship between the nuclear YAP activity and the expression of TE markers such as CDX2. Live imaging could help define the precise timeline over which YAP acts a lineage determinant and lead to further understanding of the transition of YAP functions in early embryos. In addition, the deep imaging capability provided by this reporter can illuminate previously unknown YAP activity status in embryos or other three-dimensional model systems such as organoids,. In addition, our validation of a knock-in fusion design that maintains the normal functions of endogenous YAP can serve as the ground-plan for developing other powerful genetic tools such as degrons and optogenetic tools for further functional interrogation of YAP signaling *in vivo*. In summary, we present the first KI fusion reporter mouse model to enable the readout of YAP nuclear-cytoplasmic localization by live imaging. Using this line, we reveal new aspects of dynamic YAP behaviour in preimplantation mouse embryos and the capacity to live-image postimplantation embryos with penetration at depths of up to 200μm. This live imaging YAP reporter can be combined with other appropriate reporters to study multiple developmental and disease processes.

## Acknowledgement

This work was funded by CIHR (FDN-143334) to J.R CIHR (MOP 168992) to S.H and Y.S, and Medicine by Design Grand Questions Program (Canada First Research Excellence Fund) to S.H and Y.S. The authors acknowledge technical support from the Model Production Core staff led by M. Gertsenstein at the Centre for Phenogenomics; E. Posfai (Princeton University) for early collaboration efforts on generating an N-terminal Halo-tagged YAP reporter; V. Verkhusha (Albert Einstein College of Medicine) for providing a construct encoding the emiRFP670 fluorescent protein and discussions on selecting NIR fluorescent proteins.

## Data Availability Statement

The raw data that support the findings of this study will be deposited to public databases or from the corresponding authors on reasonable request.

## Author contributions

B.G., and J.R. conceived the study. B.G. designed and produced the C-YAP reporter mouse line. B.G, B.B and M.Z designed, carried out and analyzed all experiments. J.R., Y.S and S.H. provided supervision and funding for the study. B.G and J.R wrote the manuscript and all authors reviewed and approved the manuscript.

---

**Extended Figure 1:**
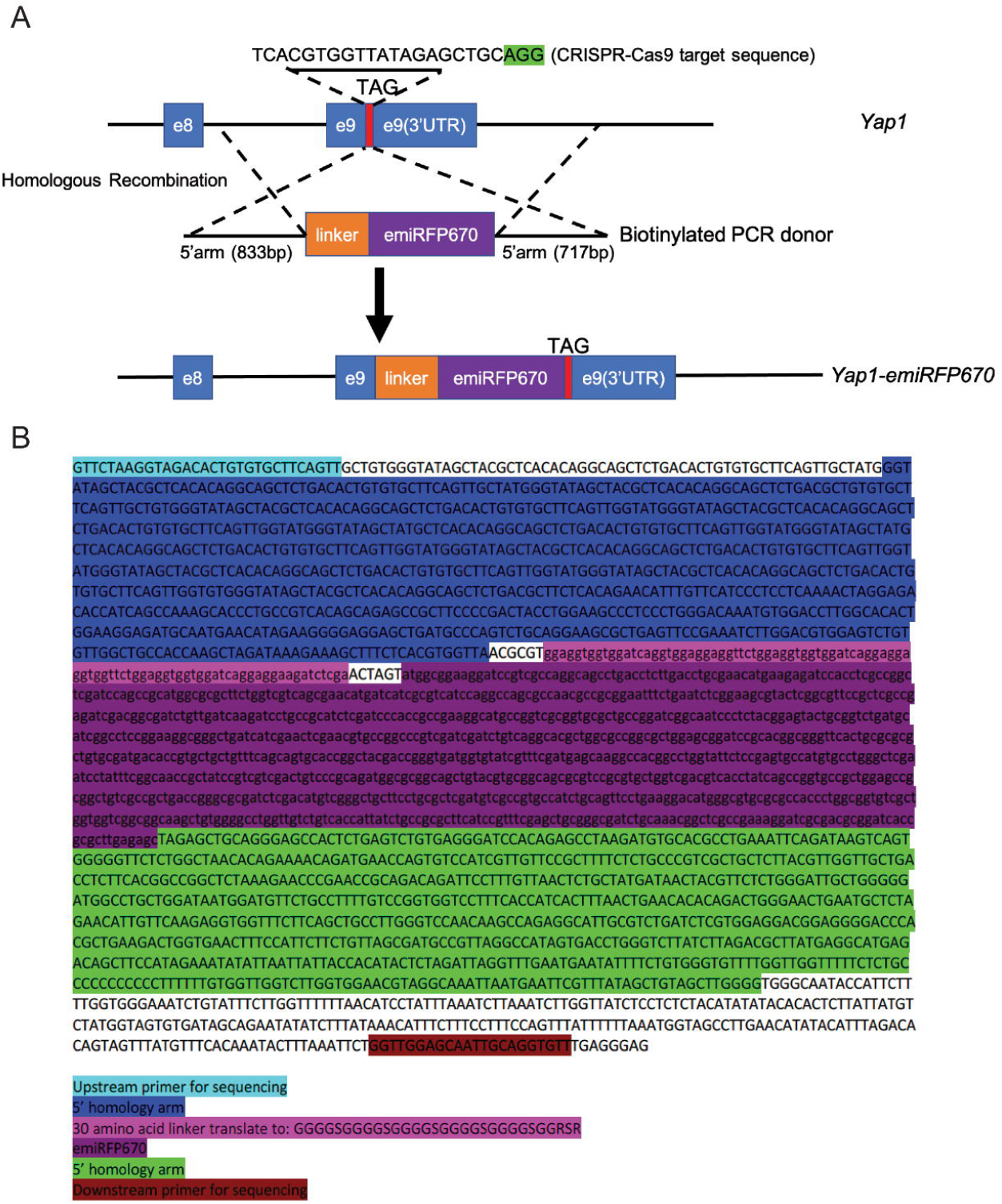
Design and sequence of the YAP-emiRFP670 allele. A. Sequence of the guide RNA and structure of the repair donor. B. Exact sequence of the KI insert and homology arms confirmed by Sanger sequencing.

**Extended Figure 2:**
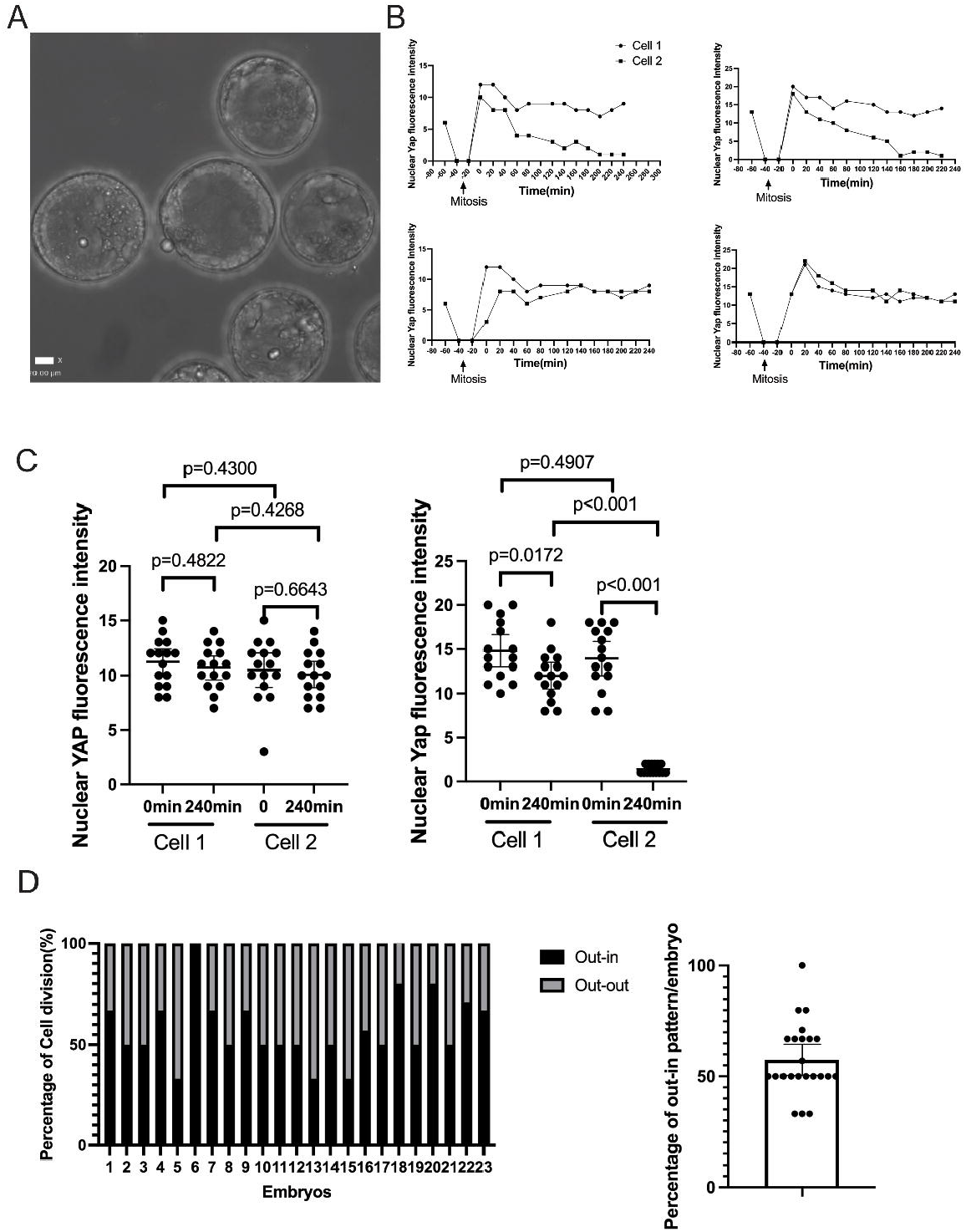
Supplementary to the live imaging analysis of 8-16 cell transition of mouse embryos. A. Viable blastocysts after a live imaging session. B. Examples of the trends of nuclear YAP fluorescence intensity after cell division during live imaging. C. Statistical analysis of nuclear YAP fluorescence intensity in each mitotic pair right after cell division (0 min) and after 240 min (Each dot represents a cell, n=15. Bars present mean and 95% confident interval (CI). Left, mitotic pairs that present out-out pattern. Right: mitotic pairs that present out-in pattern. D. Numeric analysis of cell division pattern within each embryo. Left, percentage of out-in and out-out patterned mitotic pairs in each embryo(n=23). Right, statistical analysis of percentage out-in pattern among all analyzed embryos(n=23). Presented as Mean and 95% CI. Actual mean=57.48%, one-sample t-test with the theoretical mean =50%, p=0.0381.

## Movie legends

Movie 1. Time lapse imaging of ROCKi treated YAP-emiRFP670 reporter embryos. Embryos were treated with 20μM Y-27632 and live imaged for 20 hours at a frame rate of 90min/frame.

Movie 2. Movie 2 Time lapse imaging YAP-emiRFP670 embryos from 8-16 cell stage. 8-cell stage YAP-emiRFP670 homozygous embryos were live imaged for 20 hours at a frame rate of 20min/frame.

Movie 2. Time lapse imaging YAP-emiRFP670 embryos from 8-16 cell stage. 8-cell stage YAP-emiRFP670 homozygous embryos were live imaged for 20 hours at a frame rate of 20min/frame.

Movie 3. Zoomed-in views of time lapse imaging of a single YAP-emiRFP670 embryos from 8-16 cell stage.

Movie 4. Time lapse imaging YAP-emiRFP670 embryos from 4-16 cell stage. Late 4-cell stage YAP-emiRFP670 homozygous embryos were live imaged for 28 hours at a frame rate of 20min/frame.

Movie 5. Movie 5 Zoomed in views of time lapse imaging of a single YAP-emiRFP670 embryo treated with ROCK inhibitor, illustrating that the initial YAP nuclear translocation after cell division was not affected by ROCK inhibition.

Movie 6. Rotational views of a live E8.0 (before turning) YAP-emiRFP670 embryo Movie 7. Rotational views of a live E8.5 (after turning-10-somites stage) YAP-emiRFP670 embryo.

Movie 8. Rotational views of a live E9.5 YAP-emiRFP670 embryo.

Movie 9. Movie 9 Z-stack views of the head region of a E8.5 YAP-emiRFP670 embryo at 10-somites stage.

Movie 10. Time lapse imaging of the head region of a E8.5 YAP-emiRFP670 embryo at around 10-somites stage, illustrating a population of migrating cells that presented strong nuclear YAP signaling. Left. Movie without track annotations. Right. Movie with track annotations.

Movie 11. Time-lapse imaging of preimplantation embryos with YAP-emiRFP670 and CDX2-eGFP (8-cell to 16-cell stage).

Movie 12. Time-lapse imaging of preimplantation embryos with YAP-emiRFP670 and CDX2-eGFP-zoomed in embryo 1(8-cell to 16-cell stage).

Movie 13. Time-lapse imaging of preimplantation embryos with YAP-emiRFP670 and CDX2-eGFP-zoomed in embryo 2(8-cell to 16-cell stage).

Movie 14. Time-lapse imaging of preimplantation embryos with YAP-emiRFP670 and CDX2-eGFP under ROCKi treatment.

